# Pitfalls in estimating the global carbon removal via forest expansion – a comment on Bastin et al. (2019)

**DOI:** 10.1101/788026

**Authors:** Andreas Krause, Almut Arneth, Anita Bayer, Allan Buras, Thomas Knoke, Christian Zang, Anja Rammig

## Abstract

We believe the carbon removal potential of 205 Gt C reported by Bastin et al. (2019) to be overestimated. The authors did not consider the carbon already stored on the land with identified tree restoration potential. For instance, grasslands and degraded forests have similar soil carbon stocks as old-growth forests. Accounting for these inconsistencies in the calculation, we estimate a carbon removal potential of 81 Gt C, i.e. only 39% of the Bastin et al. estimate. In addition, some of the assumptions about potential restoration area seem questionable and an essential question remains unanswered: why do areas identified as suitable for tree growth currently lack tree cover?

## Main

The potential of CO_2_ removal from the atmosphere via forest expansion is discussed since a few decades (House, Prentice, & Le Quere, 2002). Bastin et al. (2019) contribute to this discussion by combining high resolution satellite information with machine-learning to estimate an upper boundary of global tree restoration area. The approach on its own is a valuable contribution to the climate change debate as the IPCC Special Report on 1.5°C clarified the need of such a feasibility study. However, we argue that simply planting a lot of trees cannot balance fossil fuel emission to the extent claimed by Bastin et al. In direct response to their study, several scientists already highlighted some of the pitfalls with this simple approach like neglected albedo changes (Alkama & Cescatti, 2016), carbon cycle feedbacks (Jones et al., 2016), or the permanence of carbon storage. Despite the hardly disputed large uncertainties in forest carbon storage (Pan et al., 2011), the calculated carbon uptake potential of 205 Gt C (752 Gt CO_2_) is taken as quasi certain (±3% reported in their Table S2). We want to point towards inconsistencies and uncertainties in the underlying calculation and associated assumptions. We believe that the actual carbon uptake potential by tree restoration in the identified areas is substantially lower because a) the assumed per-area carbon uptake is too large; b) land is needed for food production; c) trees might not grow due to environmental constraints; or d) tress might grow anyways even without planting.

## Implausible per-area carbon uptake

Bastin et al. claim that the Earth’s surface could provide 900 ± 27 Mha additional tree canopy cover excluding existing forests, croplands and urban areas. However, even if we assume that this area is available for forest restoration, the estimated carbon uptake is based on a flawed calculation. While the main text gives the impression that it is all about vegetation carbon *(“vegetation in the potential restoration areas could store an additional 205 gigatonnes of carbon”),* the supplement clarifies that soil carbon is included as well. According to the supplement, the authors estimated carbon uptake by tree restoration by multiplying the restoration area by biome-specific total carbon densities of intact forests taken from Pan et al. (2011) and Grace, San Jose, Meir, Miranda, and Montes (2006). However, applying the carbon density of savannas (202.4 MgC ha^−1^) from Grace et al. (2006) also to tundra, deserts, montane grasslands, and Mediterranean forests is certainly a too simplistic view not acknowledging crucial features of and differences between these ecosystems. Even more worrying, the authors simply assume an initial carbon density of zero in the lands identified for tree restoration. This is unrealistic. In fact, grasslands have similar, often even larger soil carbon stocks than forests (Guo & Gifford, 2002; Li et al., 2018), whose fate after reforestation is highly uncertain in sign and magnitude (Krause et al., 2018). Comparable soil carbon stocks have also been reported for degraded forests and undisturbed forests (Chiti, Diaz-Pines, Butterbach-Bahl, Marzaioli, & Valentini, 2018; Houghton & Hackler, 2003). Adequately accounting for soil carbon changes is particularly important in the boreal zone where much of the calculated restoration area is located: Pan et al. (2011) explicitly point out that “*boreal forests have only 20% in biomass”*, meaning that in these regions the calculated carbon uptake potential might be overestimated by a factor of 5 because soil carbon storage will remain relatively constant following tree planting. The authors could have multiplied their calculated boreal forest potential by 0.2 (or 0.26, including deadwood (Pan et al., 2011)) to get a better estimate. Similar calculations can be done for temperate and tropical forests (0.39 and 0.58, respectively). Grace et al. (2006) report an even smaller vegetation fraction of 0.14 for savannas. When we apply these realistic carbon densities accounting only for vegetation carbon uptake, the global potential is reduced from 205 Gt C to 81 Gt C (Fig. 1), equivalent to only 7 years of present-day anthropogenic emissions (Le Quere et al., 2018). Interestingly, this number is similar to the 72-76 Gt C from an independent estimate (Taylor & Marconi, 2019). But even under the assumption that forests store much more soil carbon than the previous land cover, soil carbon accumulation in high latitudes would take several centuries (Krause, Pugh, Bayer, Lindeskog, & Arneth, 2016) rather than decades as stated in Bastin et al.

**Figure 1:**
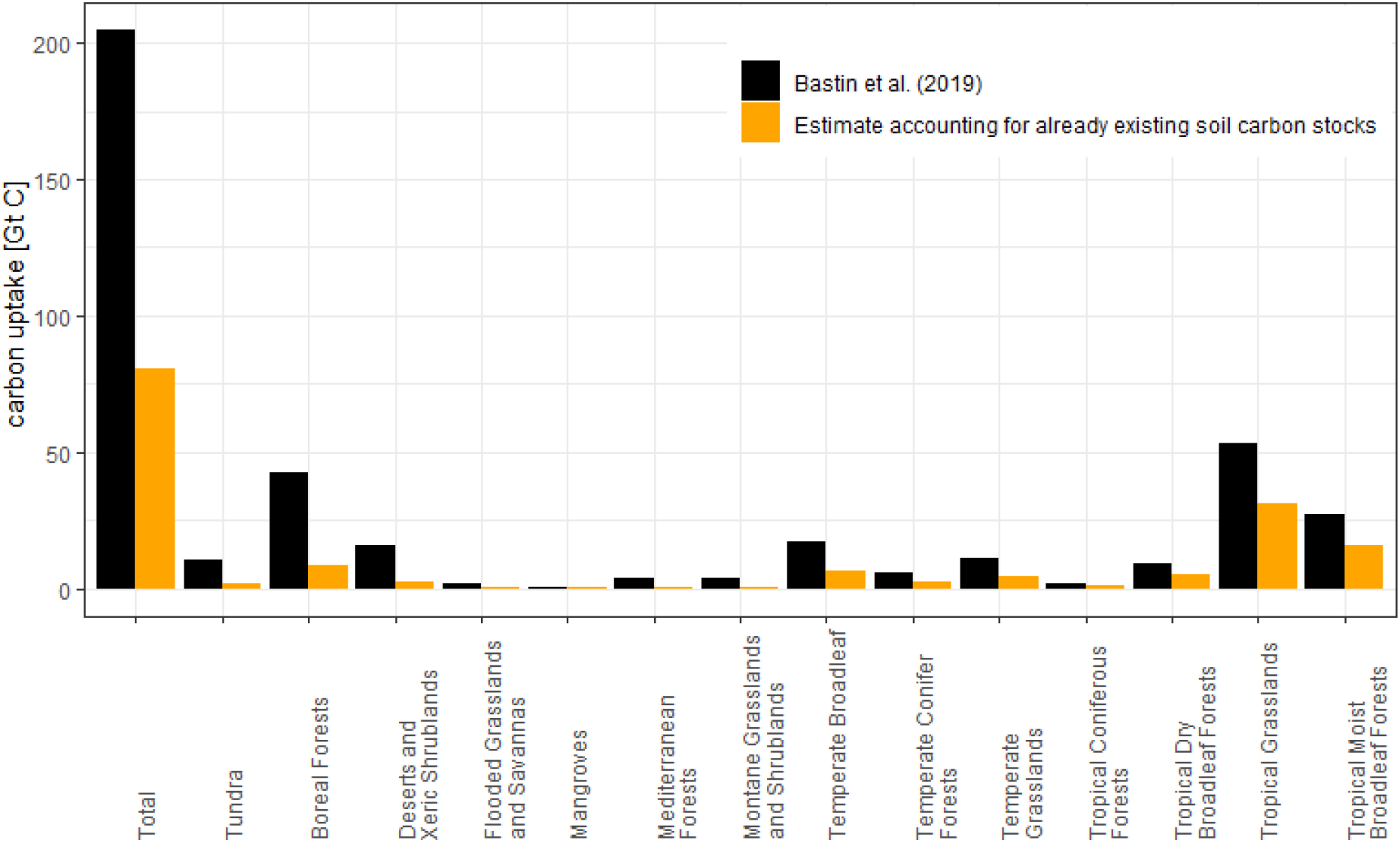
Biome-specific carbon uptake from Bastin et al. (2019) and what we consider a more realistic estimate accounting for already available soil carbon stocks. Bastin et al. did not account for the carbon currently stored on the land while we assume that forests have the same soil carbon density as the replaced vegetation, which is a much more realistic assumption (e.g. Guo & Gifford, 2002) that still doesn’t consider differences in soil carbon stocks between tree-covered and tree-free areas of the same ecosystem. Both estimates use average biome-specific total carbon densities from Pan et al. (2011) and Grace et al. (2006) and a global restoration area of 900 Mha which is likely an overestimation (see discussion below). For our estimate we multiplied the Bastin et al. values by biome-specific vegetation carbon to total carbon ratios (Grace et al., 2006; Pan et al., 2011) following the mapping of Bastin et al.: 0.2 for boreal forests, 0.39 for temperate broadleaf and conifer forests and temperate grasslands, 0.14 for tundra, deserts, Mediterranean forests, montane grasslands, flooded grasslands, and savannas (“dryland”), and 0.58 for all other biomes.

## Overestimated restoration area

In addition, the authors are very vague about what these lands identified for tree restoration actually are. There must be a reason why trees are absent from these landscapes while, according to their model, site conditions are suitable for tree growth. One possibility is that human activities have reduced tree cover below its potential maximum, in which case the question arises if the existing human interventions could be stopped and how that could be achieved. The paper states *“We kept grazing areas, as several studies suggest alternatives to improve the efficiency of livestock production”.* This is a weak justification to assume all suitable pastures to be available for restoration, given global population growth and current diet shifts towards higher per capita meat consumption. In replies to critical comments on their paper, the authors now argue that they mean to plant only two to three trees on most fields (Crown Publications, 2019; field area remains unclear, however) and that they only calculated a theoretical potential (RealClimate, 2019) but that’s not clear from the paper. Croplands were excluded from restoration areas but it seems unlikely that future population can be fed from existing cropland, at least without other detrimental side effects like water withdrawal for irrigation or increased nitrogen losses from fertilizers. The potential restoration areas could also include recently harvested forests but in this case the forest would regrow over time anyways, i.e. there is a balance between regrowth and harvest. Preventing wood harvest might increase ecosystem carbon storage but the overall carbon mitigation might be negative because of less carbon in wood products and associated substitution effects (Klein, Hollerl, Blaschke, & Schulz, 2013). Forests are also prone to disturbances like windthrow, insect outbreaks, damage caused by game animals, or wildfires (e.g. compare restoration area with Fig. 2 in Veldman et al., 2015). Additional environmental drivers not accounted for in the model presented by Bastin et al. could also prevent tree growth (e.g. soil nutrients, permafrost). In high latitudes, climate might only recently have become suitable to sustain tree growth in which case carbon accumulation would occur even without planting (Pugh et al., 2018).

We acknowledge the potential contribution of reforestation and particularly avoided tropical deforestation to climate change mitigation. However, as shown here and elsewhere, flawed calculations and neglected processes challenge the estimated carbon removal potential and derived conclusions of the Bastin et al. study. The authors themselves admit the uncertain fate of forests under future climate change, increasing disturbance rates, and atmospheric CO_2_. In addition to the issues raised here, such uncertainties also need to be considered. Clearly forest restoration can only be part of a much broader range of mitigation activities and the most effective climate change solution to date is to reduce fossil fuel emissions immediately.

